# Ecosystem’s Oxygen Deficiency and Selection of Antimicrobial Resistance Genes in a One Health Perspective

**DOI:** 10.1101/2024.05.17.594675

**Authors:** Laura Ortiz-Miravalles, Amalia Prieto, Nicolas Kieffer, Ester Vergara, Rafael Cantón, Álvaro San Millán, Fernando Baquero, Alberto Hipólito, José Antonio Escudero

## Abstract

Bacteria must face and adapt to a variety of physicochemical conditions in the environment and during infection. A key condition is the concentration of dissolved oxygen, proportional to the partial pressure of oxygen (PO_2_), which is extremely variable among environmental biogeographical areas and also compartments of the human and animal body. Here, we sought to understand if the phenotype of resistance determinants commonly found in Enterobacterales can be influenced by oxygen pressure. To do so, we have compared the MIC in aerobic and anaerobic conditions of isogenic *Escherichia coli* strains containing 136 different resistance genes against 9 antibiotic families. Our results show a complex landscape of changes in the performance of resistance genes in anaerobiosis. Certain changes are especially relevant for their intensity and the importance of the antibiotic family, like the large decreases in resistance observed against ertapenem and fosfomycin among *bla*_VIM_ ß-lactamases and certain *fos* genes, respectively; however, the *bla*_OXA-48_ ß-lactamase from the clinically relevant pOXA-48 plasmid conferred 4-fold higher ertapenem resistance in anaerobiosis. Strong changes in resistance patterns in anaerobiosis were also conserved in *Klebsiella pneumoniae*. Our results suggest that anaerobiosis is a relevant aspect that can affect the action and selective power of antibiotics for specific AMRs in different environments.

---

Antimicrobial resistance (AMR) is one of the major health concerns of the 21^st^ century globally, according to the World Health Organization (WHO)^1,2^. The dissemination of AMR occurs in the global human-animal-environmental continuum, as considered in the One Health perspective^3,4^, yet the physicochemical conditions of these different compartments are diverse. In humans and animals AMR frequency is mostly determined by antibiotic consumption, following selection of bacteria harboring antibiotic resistance genes (ARGs), frequently harbored in mobile genetic platforms such as plasmids and integron cassettes (ARCs)^5,6^. Additionally, growing antibiotic pollution in the environment also exerts a significant selective pressure on AMR bacteria^7^. The environmental pharmacokinetics of antibiotics remain poorly explored, and the laboratory conditions in which the susceptibility of pathogenic bacteria is tested (based on MIC, minimal inhibitory concentration), strongly differ from the variable ecological compartments, in animal body sites, or in the environment at large^8^.

One of these unmet conditions in laboratory tests is the variability of oxygen concentrations. Here, we sought to understand if the detoxifying performance of ARGs commonly found in Enterobacterales can be influenced by dissolved oxygen (directly proportional to the PO2, the partial pressure of oxygen) in such a way that it contributes in various degrees to the selection of AMR in aerobic/anaerobic conditions.

To investigate the impact of anaerobiosis on resistance genes, we used mainly our pMBA collection^9^ that is composed of 136 isogenic strains of *E. coli* MG1655, each containing a particular ARC as the first cassette of a class 1 integron borne by the pMBA plasmid vector^10^. The collection contains resistance genes against 9 antimicrobial families covering a broad variety of mechanisms. We have also included other clinically relevant AMR plasmids and genes. We have determined the MIC levels of every member in the collection in the presence and absence of oxygen (Figs. 1B and 1C). As a control, a strain containing the empty pMBA vector was included. To rid our analysis of the potential effects of anaerobiosis on the bacterium’s resistance -and not in the gene’s function-the *performance* of an AMR gene is measured as the fold-increase in resistance it confers compared to the empty pMBA in a given environmental condition. Additionally, the strain containing pMBA reproduced the effect of anaerobiosis found in *E. coli* without pMBA, allowing to rule out confounding effects of the vector (Fig. 1A). Our data show that several ARCs varied their antimicrobial resistance effect according to oxygen availability. For example, the aminoglycoside resistance cassette *aacA56* (Figs. 1B and 1C) increases 8-fold the MIC to amikacin in the absence of oxygen, enhancing the effect of energy-dependent antibiotic reduced uptake. In addition, some families of ARCs showed opposite trends in resistance in anaerobiosis: different variants of the metallo-ß-lactamase *bla*_VIM_ and *bla*_IMP-2_ were less efficient in anaerobiosis, reducing their performance against ertapenem between 8 and 64-fold on average (Figs. 1B and 1C). A similar decrease in resistance was observed for several fosfomycin resistance genes (*fos*) in anaerobiosis -with the exception of *fosE*, that showed a 4-fold increase (Figs. 1B). Because both *bla*_VIM_ and *fos* encode metalloenzymes (that use respectively Zn^2+^ and Mn^2+^ as cofactors) it is tempting to speculate that metal coordination is being affected in anaerobiosis.

**Figure 1.**
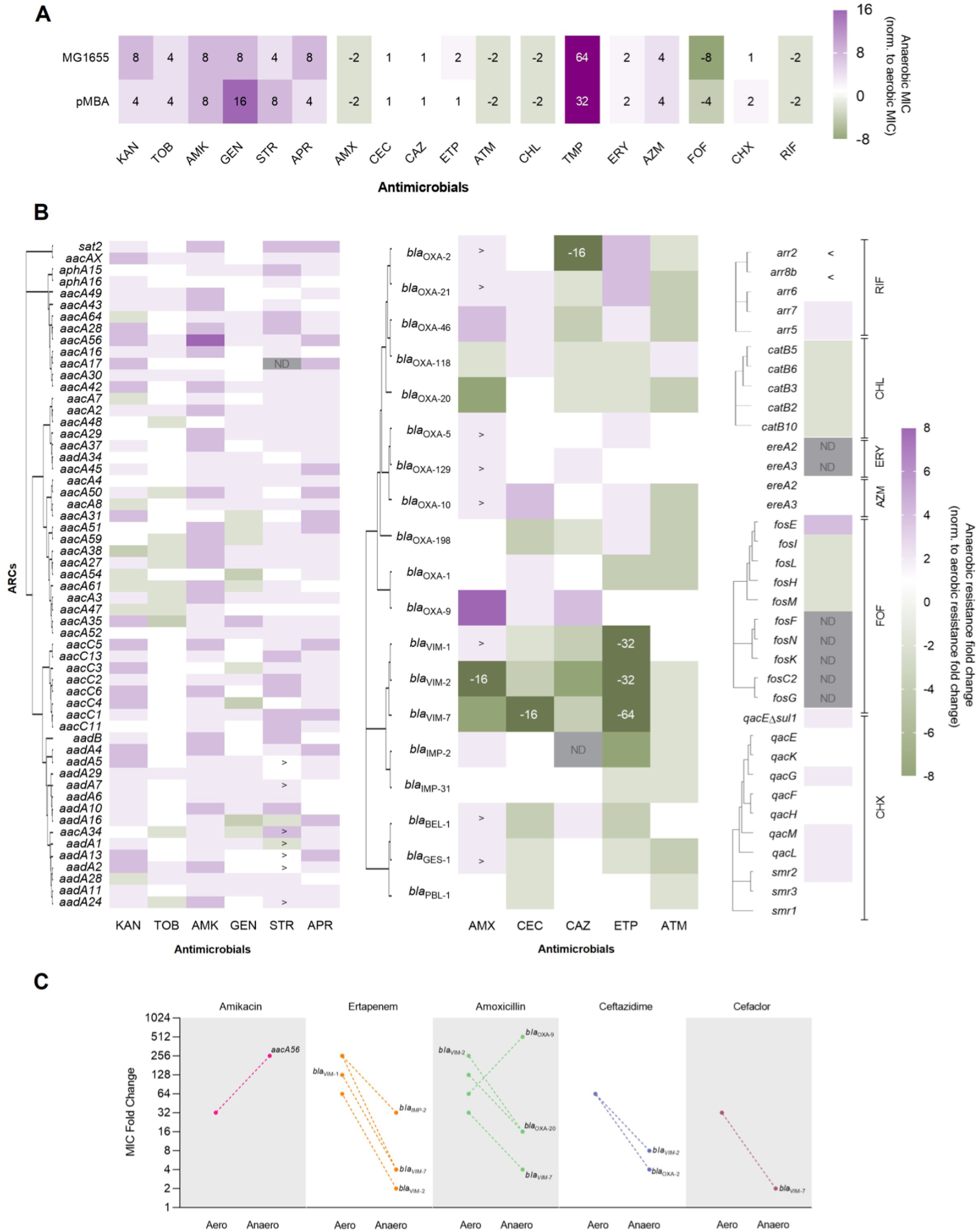
Anaerobiosis affects ARC’s function in *E. coli*. **A)** Heatmap showing the fold-change in resistance of *E. coli* MG1655 -with and without pMBA-between anaerobic and aerobic conditions against 18 antimicrobials (KAN: kanamycin, TOB: tobramycin, AMK: amikacin, GEN: gentamicin, STR: streptomycin, APR: apramycin, AMX: amoxicillin, CEC: cefaclor, CAZ: ceftazidime, ETP: ertapenem, ATM: aztreonam, CHL: chloramphenicol, TMP: trimethoprim, ERY: erythromycin, AZM: azithromycin, FOF: fosfomycin, CHX: chlorhexidine, RIF: rifampicin). Numbers in each square indicate the fold-change in resistance between anaerobic and aerobic conditions. **B)** Heatmap showing the resistance profile of each ARC in anaerobiosis, normalized by its effect in aerobic conditions. Purple indicates that an ARC confers a higher resistance increase (compared to pMBA) in anaerobic conditions than in aerobic conditions, while green represents a lower resistance increase (compared to pMBA) in anaerobiosis compared to aerobiosis. Red squares depict cases for which a precise MIC could not be determined (strains showing growth at the highest concentration of antibiotic tested) in any of both conditions. Symbols > and < are used to indicate strains that grew at the highest concentration of antibiotic tested in anaerobic or aerobic conditions respectively. A hierarchical clustering tree showing protein sequence similarity is shown in the left side of each panel for (from left to right) aminoglycoside resistance cassettes, ß-lactam resistance cassettes, other ARCs. **C)** MIC fold change (compared to pMBA) in aerobiosis and anaerobiosis of the subset of ARCs, showing at least an 8-fold difference between conditions. Points represent the mode of at least three biological replicates.

At least 8 ARCs showed strong changes in their performance in anaerobiosis (defined as fold changes ≥ 8-fold between conditions) (Fig. 1C). This can be of clinical relevance, as carbapenemases inactivating last resource antimicrobial agents, such as *bla*_VIMs_ and *bla*_IMP-2_, confer lower resistance levels in anaerobiosis. Our data suggests that bacteria carrying these metallo-ß-lactamase genes could become susceptible to drugs such as ertapenem in anaerobic compartments of the animal or human body (such as the large intestine), including infection sites or in particular environmental spaces. Yet, the conventional antibiogram would yield an ertapenem resistance phenotype, wrongly preventing clinicians from using a possibly successful drug. The opposite trend is observed for other ß-lactamases, such as OXA-9 and OXA-48 ß-lactamases. Indeed, plasmid pOXA-48 confers resistance to ertapenem in *E. coli* under anaerobic conditions but the strain is considered susceptible to this antibiotic under aerobic conditions. A similar phenomenon is observed for plasmid R388^11^ containing *bla*_VEB-1_, where resistance to ß-lactam antibiotic ceftazidime increases in anaerobiosis (Fig. 2A). The resistance profile of an *mcr-1* did not show relevant changes (≥ 4-fold change) in anaerobic conditions (Fig 2A).

**Figure 2.**
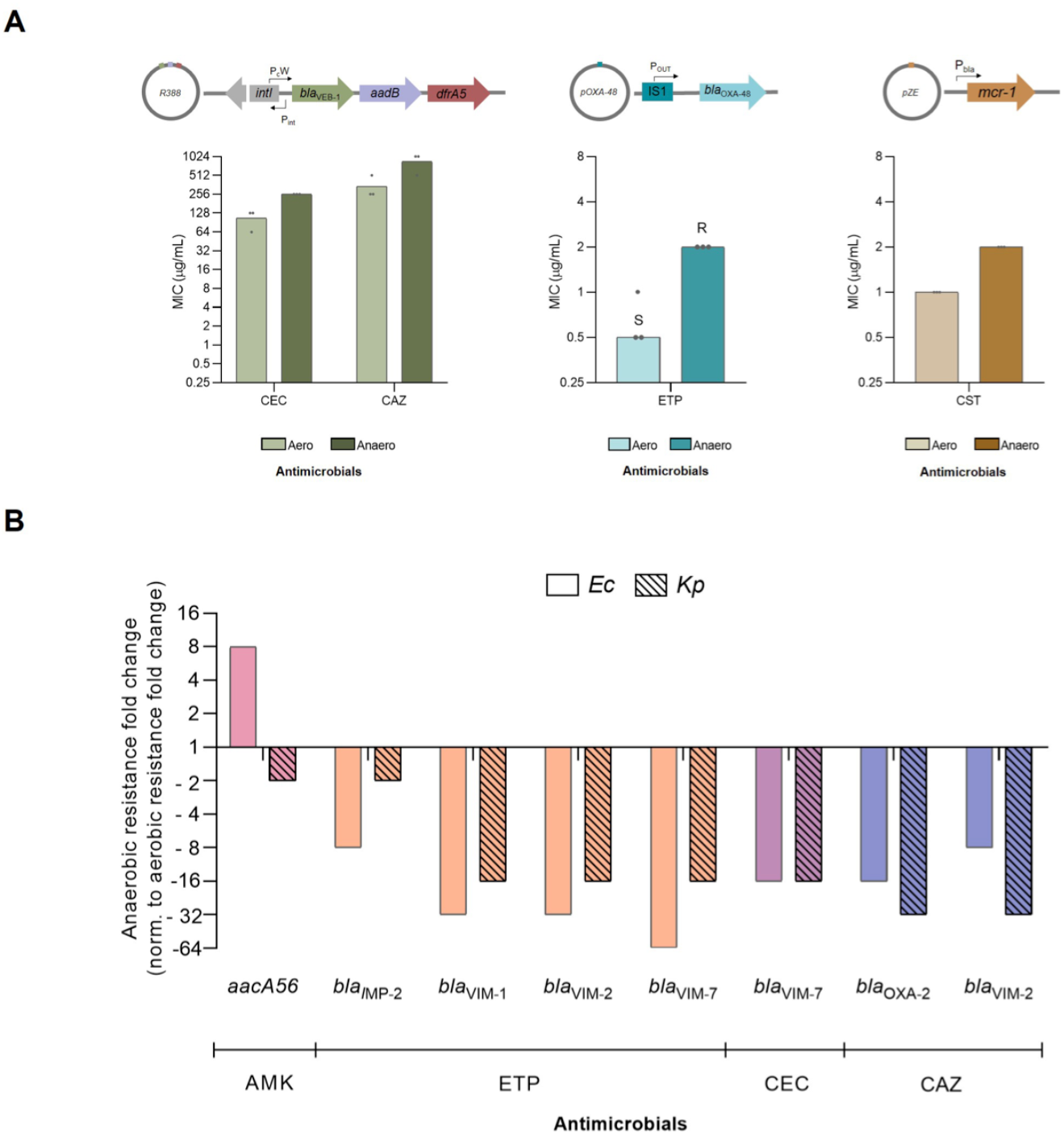
Anaerobiosis modulates antimicrobial susceptibility across different species. **A)** MIC of *E. coli* MG1655 harboring clinically relevant resistance genes and plasmids, against several antimicrobials. Bars represent the mode of three biological replicates (black dots). Light and dark colored bars represent aerobic and anaerobic measures respectively. S and R letters stand for susceptible and resistant according to EUCAST clinical breakpoints. A scheme illustrating the genetic background of each ARG is displayed above each graph. **B)** Comparison between anaerobic MIC fold change normalized to aerobic MIC fold change of pMBA_ARC_-containing *E. coli* and *K. pneumoniae*. The effect of oxygen concentration on ARCs is similar in both species. MIC fold change is calculated as the mode of at least six biological replicates. Non-patterned and patterned columns represent data retrieved from *E. coli* and *K. pneumoniae* respectively.

To test if these results are conserved in other Enterobacterales, we selected the ARCs showing ≥ 8-fold change between conditions in *E. coli* and introduced them in strain ATCC23357 of *Klebsiella pneumoniae*. The trends in resistance changes were similar between both species (Fig. 2B).

In this work we show how oxygen availability, as an example of physico-chemical conditions of ecosystems can influence AMR. Our results have implications in clinical treatments but also in environmental settings. Indeed, in natural water bodies the amount of dissolved oxygen depends on the depth of the water column and the action of winds, waves, photosynthetic organisms, temperature, and salinity. Water bodies are anthropogenically polluted with antibiotics and antibiotic-resistant bacteria, many of them originating from human and animal microbiota^12^. Hence, oxygen availability can influence selection of AMR organisms in the environment, in particular in sewage and sludge water treatment plants, that are frequently based on treatments decreasing dissolved oxygen^13^. Altogether, this means that the possibility of modifying the effect on the AMR genes is ecologically and geographically dependent, and AMR surveillance research on the relation of oxygen and the abundance of AMRs should probably be developed in the near future^14^. Finally, we should mention the consequences of global warming, increasing bacterial growth but reducing the availability of dissolved oxygen in different biogeographical scenarios, thus altering the environmental selection of AMR^15^.

## Data Availability

Data sharing not applicable to this article as no datasets other than what is shown in figures were generated or analysed during the current study.

## Acknowledgements

We would like to thank the members of MBA lab for their helpful comments and the funding agencies for their support.

