## Supplementary material for "Ecosystem’s Oxygen Deficiency and Selection of Antimicrobial Resistance Genes in a One Health Perspective": supp

### Supplementary Table S1. Clinical strains used in this study.

| Name | Specie | Plasmid content | Resistance genes | Source |
|---|---|---|---|---|
| EC05 | *E. coli* | Col-like, IncFII, IncI | *aadA5, blaCTX-M-1, sul2, tet(A), mdf(A)* | R-GNOSIS project |
| EC10 | *E. coli* | Col-like, IncFII, pEC4115 | *aac(3)-Iva, aph(3'')-Ib, aph(4)-Ia, aph(6)-Id, blaCTX-M-1, mph(A), mdf(A)* | R-GNOSIS project |
| EC14 | *E. coli* | Col-like, IncFIB, IncFII | *aac(3)-Iid, aadA2, blaCTX-M-32, cmlA1, sul3, tet(B), mdf(A)* | R-GNOSIS project |
| Kpn04 | *K. pneumoniae* | IncFIB, IncFII, IncX, IncFIA, IncHI1B | *aac(3)-Iia, aac(6')-Ib-cr, aph(3'')-Ib, aph(6)-Id, blaCTX-M-15, blaOXA-1, blaSHV-187, blaTEM-1B, dfrA14, fosA, qnrB1, sul2, tet(A), oqxA, oqxB* | R-GNOSIS project |
| Kpn13 | *K. pneumoniae* | Col-like, IncFII, IncFIA, IncHI1B | *aph(3'')-Ib, aph(6)-Id, blaCTX-M-15, blaSHV-187, blaTEM-1B, dfrA14, fosA, qnrB1, sul2, oqxA, oqxB* | R-GNOSIS project |
| Kpn21 | *K. pneumoniae* | Col-like, IncFIB | *aadA2, aph(3'')-Ib, aph(6)-Id, blaCTX-M-15, blaSHV-60, catA2, dfrA12, fosA, mph(A), qnrS1, sul1, sul2, tet(A), oqxA, oqxB* | R-GNOSIS project |

### Supplementary Table S2. Strains and plasmids used in this study.

All strains constructed for this work were built in an *E. coli* MG1655 or *K. pneumoniae* ATCC23357 backgrounds (references A072 and B929 in lab collection).

| Strain | Genetic background | Plasmid | ARC cloned | Source |
|---|---|---|---|---|
| A390 | MG1655 | pMBA $_{\textit{bla}\text{BEL-1}}$ | *bla*$_{\text{BEL-1}}$ | *Hipólito et al*, 2023 |
| A380 | MG1655 | pMBA $_{\textit{bla}\text{GES-1}}$ | *bla*$_{\text{GES-1}}$ | *Hipólito et al*, 2023 |
| A387 | MG1655 | pMBA $_{\textit{bla}\text{IMP-2}}$ | *bla*$_{\text{IMP-2}}$ | *Hipólito et al*, 2023 |
| A383 | MG1655 | pMBA $_{\textit{bla}\text{IMP-31}}$ | *bla*$_{\text{IMP-31}}$ | *Hipólito et al*, 2023 |
| A441 | MG1655 | pMBA $_{\textit{bla}\text{OXA-1}}$ | *bla*$_{\text{OXA-1}}$ | *Hipólito et al*, 2023 |
| A373 | MG1655 | pMBA $_{\textit{bla}\text{OXA-2}}$ | *bla*$_{\text{OXA-2}}$ | *Hipólito et al*, 2023 |
| A381 | MG1655 | pMBA $_{\textit{bla}\text{OXA-5}}$ | *bla*$_{\text{OXA-5}}$ | *Hipólito et al*, 2023 |
| A384 | MG1655 | pMBA $_{\textit{bla}\text{OXA-9}}$ | *bla*$_{\text{OXA-9}}$ | *Hipólito et al*, 2023 |
| A374 | MG1655 | pMBA $_{\textit{bla}\text{OXA-10}}$ | *bla*$_{\text{OXA-10}}$ | *Hipólito et al*, 2023 |
| A385 | MG1655 | pMBA $_{\textit{bla}\text{OXA-20}}$ | *bla*$_{\text{OXA-20}}$ | *Hipólito et al*, 2023 |
| A385 | MG1655 | pMBA $_{\textit{bla}\text{OXA-20}}$ | *bla*$_{\text{OXA-20}}$ | *Hipólito et al*, 2023 |
| A382 | MG1655 | pMBA $_{\textit{bla}\text{OXA-21}}$ | *bla*$_{\text{OXA-21}}$ | *Hipólito et al*, 2023 |
| A395 | MG1655 | pMBA $_{\textit{bla}\text{OXA-46}}$ | *bla*$_{\text{OXA-46}}$ | *Hipólito et al*, 2023 |
| A375 | MG1655 | pMBA $_{\textit{bla}\text{OXA-118}}$ | *bla*$_{\text{OXA-118}}$ | *Hipólito et al*, 2023 |
| A391 | MG1655 | pMBA $_{\textit{bla}\text{OXA-129}}$ | *bla*$_{\text{OXA-129}}$ | *Hipólito et al*, 2023 |
| A376 | MG1655 | pMBA $_{\textit{bla}\text{OXA-198}}$ | *bla*$_{\text{OXA-198}}$ | *Hipólito et al*, 2023 |
| A803 | MG1655 | pMBA $_{\textit{bla}\text{PBL-1}}$ | *bla*$_{\text{PBL-1}}$ | *Hipólito et al*, 2023 |
| A388 | MG1655 | pMBA $_{\textit{bla}\text{VIM-1}}$ | *bla*$_{\text{VIM-1}}$ | *Hipólito et al*, 2023 |
| A396 | MG1655 | pMBA $_{\textit{bla}\text{VIM-2}}$ | *bla*$_{\text{VIM-2}}$ | *Hipólito et al*, 2023 |
| A389 | MG1655 | pMBA $_{\textit{bla}\text{VIM-7}}$ | *bla*$_{\text{VIM-7}}$ | *Hipólito et al*, 2023 |
| A327 | MG1655 | pMBA $_{\textit{aacA2}}$ | *aacA2* | *Hipólito et al*, 2023 |
| A266 | MG1655 | pMBA $_{\textit{aacA3}}$ | *aacA3* | *Hipólito et al*, 2023 |

| | | | | |
|---|---|---|---|---|
| A328 | MG1655 | pMBA *aacA4* | *aacA4* | *Hipólito et al*, 2023 |
| A267 | MG1655 | pMBA *aacA7* | *aacA7* | *Hipólito et al*, 2023 |
| A268 | MG1655 | pMBA *aacA8* | *aacA8* | *Hipólito et al*, 2023 |
| A263 | MG1655 | pMBA *aacA16* | *aacA16* | *Hipólito et al*, 2023 |
| A543 | MG1655 | pMBA *aacA17* | *aacA17* | *Hipólito et al*, 2023 |
| A264 | MG1655 | pMBA *aacA27* | *aacA27* | *Hipólito et al*, 2023 |
| A256 | MG1655 | pMBA *aacA28* | *aacA28* | *Hipólito et al*, 2023 |
| A257 | MG1655 | pMBA *aacA29* | *aacA29* | *Hipólito et al*, 2023 |
| A270 | MG1655 | pMBA *aacA30* | *aacA30* | *Hipólito et al*, 2023 |
| B657 | MG1655 | pMBA *aacA31* | *aacA31* | *Hipólito et al*, 2023 |
| A272 | MG1655 | pMBA *aacA34* | *aacA34* | *Hipólito et al*, 2023 |
| A329 | MG1655 | pMBA *aacA35* | *aacA35* | *Hipólito et al*, 2023 |
| A258 | MG1655 | pMBA *aacA37* | *aacA37* | *Hipólito et al*, 2023 |
| A603 | MG1655 | pMBA *aacA38* | *aacA38* | *Hipólito et al*, 2023 |
| A273 | MG1655 | pMBA *aacA42* | *aacA42* | *Hipólito et al*, 2023 |
| C117 | MG1655 | pMBA *aacA43* | *aacA43* | *Hipólito et al*, 2023 |
| A265 | MG1655 | pMBA *aacA45* | *aacA45* | *Hipólito et al*, 2023 |
| A274 | MG1655 | pMBA *aacA47* | *aacA47* | *Hipólito et al*, 2023 |
| A259 | MG1655 | pMBA *aacA48* | *aacA48* | *Hipólito et al*, 2023 |
| B656 | MG1655 | pMBA *aacA49* | *aacA49* | *Hipólito et al*, 2023 |
| A260 | MG1655 | pMBA *aacA50* | *aacA50* | *Hipólito et al*, 2023 |
| A261 | MG1655 | pMBA *aacA51* | *aacA51* | *Hipólito et al*, 2023 |
| A326 | MG1655 | pMBA *aacA52* | *aacA52* | *Hipólito et al*, 2023 |
| B655 | MG1655 | pMBA *aacA54* | *aacA54* | *Hipólito et al*, 2023 |
| A262 | MG1655 | pMBA *aacA56* | *aacA56* | *Hipólito et al*, 2023 |
| A303 | MG1655 | pMBA *aacA59* | *aacA59* | *Hipólito et al*, 2023 |
| A302 | MG1655 | pMBA *aacA61* | *aacA61* | *Hipólito et al*, 2023 |
| A566 | MG1655 | pMBA *aacA64* | *aacA64* | *Hipólito et al*, 2023 |
| A331 | MG1655 | pMBA *aacAX* | *aacAX* | *Hipólito et al*, 2023 |
| A308 | MG1655 | pMBA *aacC1* | *aacC1* | *Hipólito et al*, 2023 |
| A304 | MG1655 | pMBA *aacC2* | *aacC2* | *Hipólito et al*, 2023 |
| B625 | MG1655 | pMBA *aacC3* | *aacC3* | *Hipólito et al*, 2023 |
| A320 | MG1655 | pMBA *aacC4* | *aacC4* | *Hipólito et al*, 2023 |
| A305 | MG1655 | pMBA *aacC5* | *aacC5* | *Hipólito et al*, 2023 |
| A306 | MG1655 | pMBA *aacC6* | *aacC6* | *Hipólito et al*, 2023 |
| A332 | MG1655 | pMBA *aacC11* | *aacC11* | *Hipólito et al*, 2023 |
| A309 | MG1655 | pMBA *aacC13* | *aacC13* | *Hipólito et al*, 2023 |
| A311 | MG1655 | pMBA *aadA1* | *aadA1* | *Hipólito et al*, 2023 |
| A333 | MG1655 | pMBA *aadA2* | *aadA2* | *Hipólito et al*, 2023 |
| A319 | MG1655 | pMBA *aadA4* | *aadA4* | *Hipólito et al*, 2023 |
| A313 | MG1655 | pMBA *aadA5* | *aadA5* | *Hipólito et al*, 2023 |
| A321 | MG1655 | pMBA *aadA6* | *aadA6* | *Hipólito et al*, 2023 |
| A397 | MG1655 | pMBA *aadA7* | *aadA7* | *Hipólito et al*, 2023 |
| A315 | MG1655 | pMBA *aadA10* | *aadA10* | *Hipólito et al*, 2023 |
| A312 | MG1655 | pMBA *aadA11* | *aadA11* | *Hipólito et al*, 2023 |
| A322 | MG1655 | pMBA *aadA13* | *aadA13* | *Hipólito et al*, 2023 |
| A316 | MG1655 | pMBA *aadA16* | *aadA16* | *Hipólito et al*, 2023 |
| A318 | MG1655 | pMBA *aadA24* | *aadA24* | *Hipólito et al*, 2023 |
| A604 | MG1655 | pMBA *aadA28* | *aadA28* | *Hipólito et al*, 2023 |
| A314 | MG1655 | pMBA *aadA29* | *aadA29* | *Hipólito et al*, 2023 |
| A398 | MG1655 | pMBA *aadA34* | *aadA34* | *Hipólito et al*, 2023 |

| | | | | |
|---|---|---|---|---|
| A292 | MG1655 | pMBA $_{aadB}$ | *aadB* | *Hipólito et al*, 2023 |
| A422 | MG1655 | pMBA $_{aphA15}$ | *aphA15* | *Hipólito et al*, 2023 |
| C116 | MG1655 | pMBA $_{aphA16}$ | *aphA16* | *Hipólito et al*, 2023 |
| A293 | MG1655 | pMBA $_{sat2}$ | *sat2* | *Hipólito et al*, 2023 |
| A368 | MG1655 | pMBA $_{arr2}$ | *arr2* | *Hipólito et al*, 2023 |
| A334 | MG1655 | pMBA $_{arr5}$ | *arr5* | *Hipólito et al*, 2023 |
| A363 | MG1655 | pMBA $_{arr6}$ | *arr6* | *Hipólito et al*, 2023 |
| A440 | MG1655 | pMBA $_{arr7}$ | *arr7* | *Hipólito et al*, 2023 |
| A364 | MG1655 | pMBA $_{arr8b}$ | *arr8b* | *Hipólito et al*, 2023 |
| A335 | MG1655 | pMBA $_{catB2}$ | *catB2* | *Hipólito et al*, 2023 |
| A336 | MG1655 | pMBA $_{catB3}$ | *catB3* | *Hipólito et al*, 2023 |
| A365 | MG1655 | pMBA $_{catB5}$ | *catB5* | *Hipólito et al*, 2023 |
| A625 | MG1655 | pMBA $_{catB6}$ | *catB6* | *Hipólito et al*, 2023 |
| A393 | MG1655 | pMBA $_{catB10}$ | *catB10* | *Hipólito et al*, 2023 |
| A338 | MG1655 | pMBA $_{ereA2}$ | *ereA2* | *Hipólito et al*, 2023 |
| A423 | MG1655 | pMBA $_{ereA3}$ | *ereA3* | *Hipólito et al*, 2023 |
| A657 | MG1655 | pMBA $_{fosC2}$ | *fosC2* | *Hipólito et al*, 2023 |
| A339 | MG1655 | pMBA $_{fosE}$ | *fosE* | *Hipólito et al*, 2023 |
| A624 | MG1655 | pMBA $_{fosF}$ | *fosF* | *Hipólito et al*, 2023 |
| A354 | MG1655 | pMBA $_{fosG}$ | *fosG* | *Hipólito et al*, 2023 |
| A355 | MG1655 | pMBA $_{fosH}$ | *fosH* | *Hipólito et al*, 2023 |
| A356 | MG1655 | pMBA $_{fosI}$ | *fosI* | *Hipólito et al*, 2023 |
| A626 | MG1655 | pMBA $_{fosK}$ | *fosK* | *Hipólito et al*, 2023 |
| A621 | MG1655 | pMBA $_{fosL}$ | *fosL* | *Hipólito et al*, 2023 |
| A622 | MG1655 | pMBA $_{fosM}$ | *fosM* | *Hipólito et al*, 2023 |
| A627 | MG1655 | pMBA $_{fosN}$ | *fosN* | *Hipólito et al*, 2023 |
| A359 | MG1655 | pMBA $_{smr2}$ | *smr2* | *Hipólito et al*, 2023 |
| A366 | MG1655 | pMBA $_{smr3}$ | *smr3* | *Hipólito et al*, 2023 |
| A367 | MG1655 | pMBA $_{qacE}$ | *qacE* | *Hipólito et al*, 2023 |
| B091 | MG1655 | pMBA $_{qacE\Delta\text{-}sul1}$ | *qacEΔ-sul1* | *Hipólito et al*, 2023 |
| A323 | MG1655 | pMBA $_{qacF}$ | *qacF* | *Hipólito et al*, 2023 |
| A357 | MG1655 | pMBA $_{qacG}$ | *qacG* | *Hipólito et al*, 2023 |
| A324 | MG1655 | pMBA $_{qacH}$ | *qacH* | *Hipólito et al*, 2023 |
| A623 | MG1655 | pMBA $_{qacK}$ | *qacK* | *Hipólito et al*, 2023 |
| A325 | MG1655 | pMBA $_{qacL}$ | *qacL* | *Hipólito et al*, 2023 |
| A358 | MG1655 | pMBA $_{qacM}$ | *qacM* | *Hipólito et al*, 2023 |
| A082 | MG1655 | R388 $_{bla\text{VEB-1}\text{-}aadB\text{-}dfrA5}$ | *bla*$_{\text{VEB-1}}$-*aadB-dfrA5* | *Souque et al*, 2021 |
| C965 | MG1655 | pZE $_{mcr1}$ | *mcr1* | Lab collection |
| C960 | MG1655 | pOXA48 | *bla*$_{\text{OXA-48}}$ | Lab collection |
| C408 | Kpn ATCC23357 | pMBA empty vector | - | Lab collection |
| C391 | Kpn ATCC23357 | pMBA $_{aacA56}$ | *aacA56* | Lab collection |
| C418 | Kpn ATCC23357 | pMBA $_{bla\text{IMP-2}}$ | *bla*$_{IMP\text{-}2}$ | Lab collection |
| C419 | Kpn ATCC23357 | pMBA $_{bla\text{VIM-1}}$ | *bla*$_{\text{VIM-1}}$ | Lab collection |
| C422 | Kpn ATCC23357 | pMBA $_{bla\text{VIM-2}}$ | *bla*$_{\text{VIM-2}}$ | Lab collection |
| C969 | Kpn ATCC23357 | pMBA $_{bla\text{VIM-7}}$ | *bla*$_{\text{VIM-7}}$ | Lab collection |
| C967 | Kpn ATCC23357 | pMBA $_{bla\text{OXA-2}}$ | *bla*$_{\text{OXA-2}}$ | Lab collection |

**Supplementary Table S3. Antimicrobial compounds used in this study.**

| Reference | Trader | Antimicrobial |
|---|---|---|
| A1774-1G | Sigma Aldrich | Amikacin (AMK) |
| A8523-5G | Sigma Aldrich | Amoxicillin (AMX) |
| A2024-1G | Sigma Aldrich | Apramycin sulfate salt (APR) |
| PHR1088-1G | Sigma Aldrich | Azithromycin (AZM) |
| PZ0038-25MG | Sigma Aldrich | Aztreonam (ATM) |
| C6895-1G | Sigma Aldrich | Cefaclor (CEC) |
| CDS020667-50MG | Sigma Aldrich | Ceftazidime (CAZ) |
| C0378-25G | Sigma Aldrich | Chloramphenicol (CHL) |
| 282227-1G | Sigma Aldrich | Chlorhexidine (CHX) |
| E5389-1G | Sigma Aldrich | Erythromycin (ERY) |
| 901967-H | MSD | Ertapenem (ETP). INVANZ |
| G1914-5G | Sigma Aldrich | Gentamicin sulfate (GEN) |
| K4000-5G | Sigma Aldrich | Kanamycin sulfate (KAN) |
| P5396-1G | Sigma Aldrich | Phosphomycin disodium salt (FOF) |
| R3501-1G | Sigma Aldrich | Rifampicin (RIF) |
| S6501-25G | Sigma Aldrich | Streptomycin sulfate salt (STR) |
| PHR1079-1G | Sigma Aldrich | Tobramycin (TOB) |
| T7883-5G | Sigma Aldrich | Trimethoprim (TMP) |

**Supplementary Table S4. Antibiotic discs used in this study (BioRad).**

| Reference | Antibiotic (Ab.) | Ab. content (µg) |
|---|---|---|
| 66148 | Amikacin (AMK) | 30 |
| 68042 | Amoxicillin (AMX) | 20 |
| 66178 | Amoxicillin + clavulanic (AMC) | 20+10 |
| 66928 | Aztreonam (ATM) | 30 |
| 66098 | Cefepime (FEP) | 30 |
| 66368 | Cefotaxime (CTX) | 30 |
| 66228 | Cefoxitin (FOX) | 30 |
| 66218 | Cephalotin (CEF) | 30 |
| 66278 | Chloramphenicol (CHL) | 30 |
| 68648 | Ciprofloxacin (CIP) | 5 |
| 67518 | Ertapenem (ETP) | 10 |
| 67658 | Fosfomycin (FOF) | 200 |
| 66608 | Gentamicin (GEN) | 10 |
| CT0455B | Imipinem (IMP) | 10 |
| 66618 | Kanamycin (KAN) | 30 |
| 67418 | Streptomycin (STR) | 10 |
| Prepared in house | Sulfamethoxazol (RL) | 25 |
| 67448 | Tetracycline (TET) | 30 |
| 67488 | Tobramycin (TOB) | 10 |
| Prepared in house | Trimethoprim (TMP) | 5 |
| 68898 | Trimethoprim + Sulfamethoxazole (SXT) | 25 |
| 66101 | Non impregnated paper disc | |

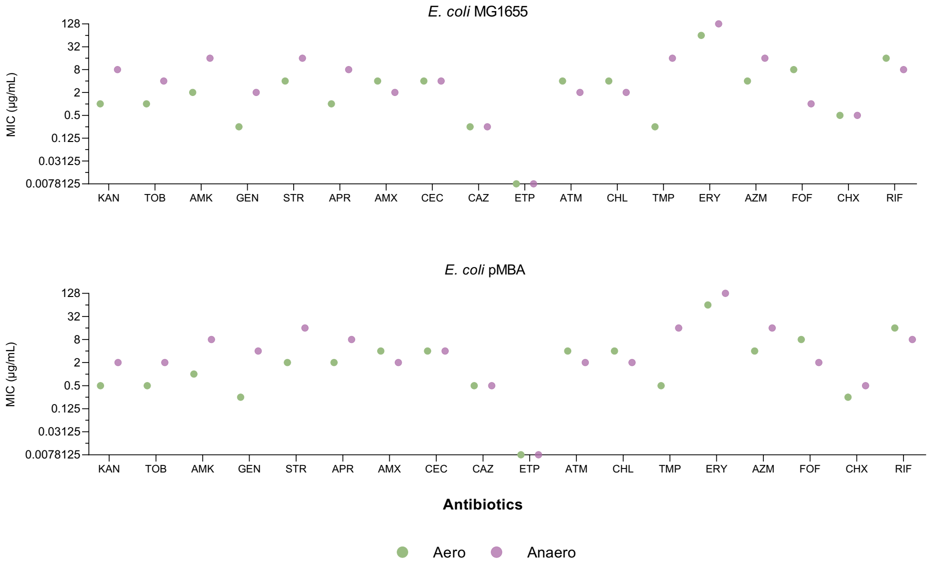


**Supplementary Figure S1. MIC quantitation for *E. coli* MG1655 and *E. coli* pMBA.** MIC values are shown as the mode of at least three biological replicates. Green dots represent aerobic measures while purple dots represent anaerobic MICs. KAN: kanamycin, TOB: tobramycin, AMK: amikacin, GEN: gentamicin, STR: streptomycin, APR: apramycin, AMX: amoxicillin, CEC: cefaclor, CAZ: ceftazidime, ETP: ertapenem, ATM: aztreonam, CHL: chloramphenicol, TMP: trimethoprim, ERY: erythromycin, AZM: azithromycin, FOF: fosfomycin, CHX: chlorhexidine, RIF: rifampicin.

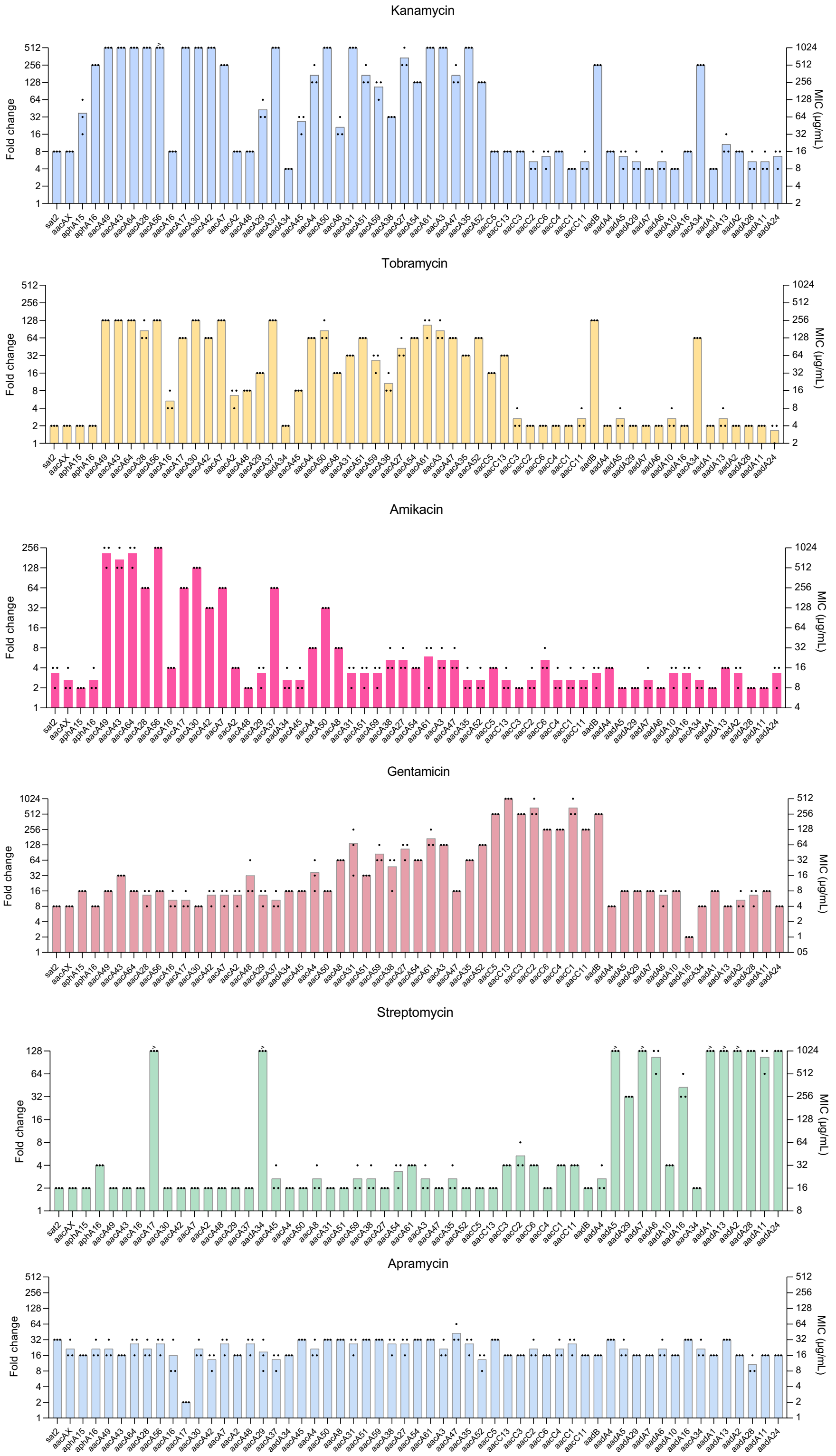


**Supplementary Figure S2. MICs of aminoglycoside resistance cassettes in anaerobiosis.** Antimicrobial resistance to kanamycin, tobramycin, amikacin, gentamicin, streptomycin, and apramycin is shown as MIC (µg/mL) in the right axis, and resistance fold increase compared to pMBA in the left axis. The MIC is the mean of three biological replicates (black dots) for each strain.

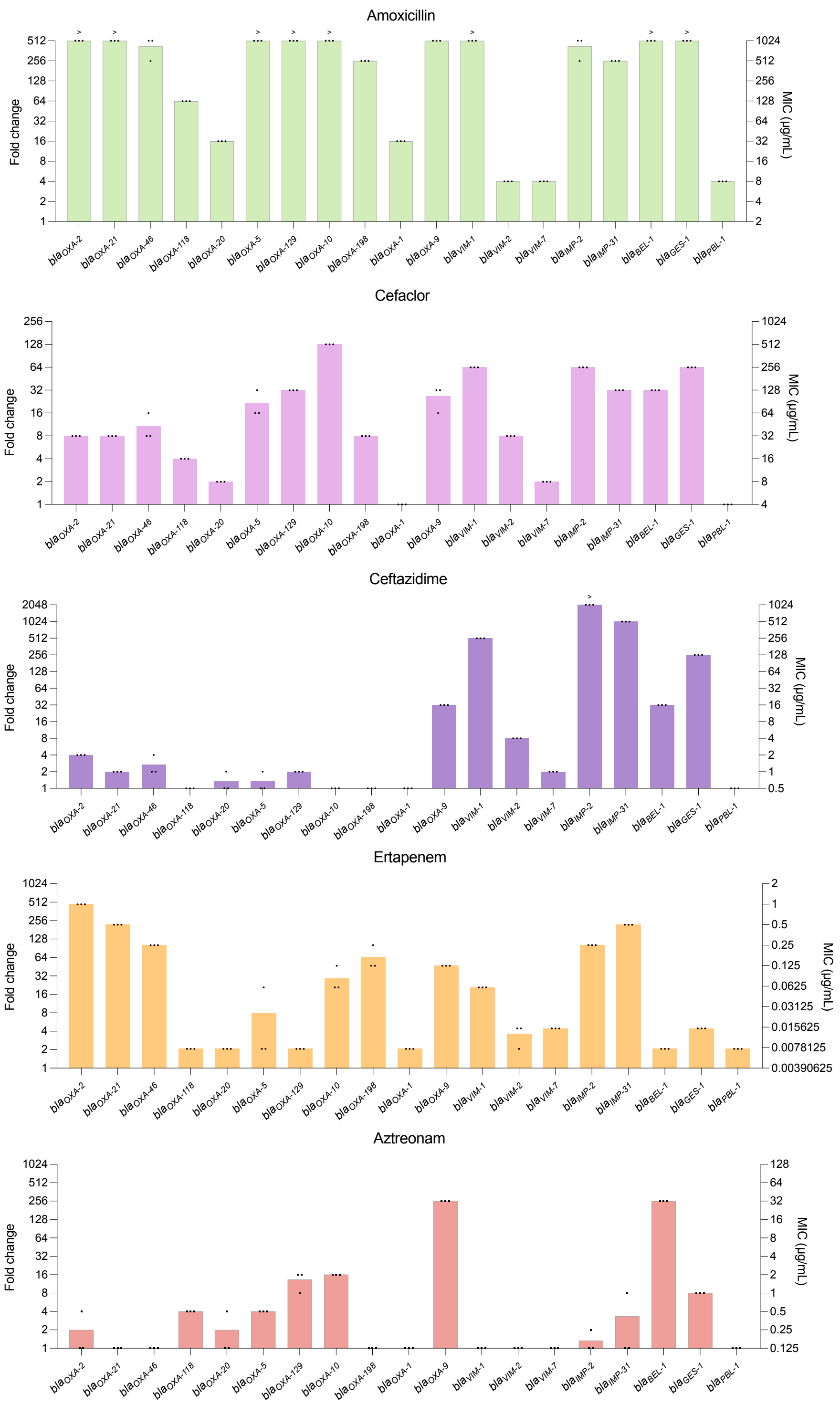


**Supplementary Figure S3. MICs of beta-lactams resistance cassettes in anaerobiosis.** Antimicrobial resistance to amoxicillin, cefaclor, ceftazidime, ertapenem, and aztreonam is shown as MIC (µg/mL) in the right axis, and resistance fold increase compared to pMBA in the left axis. The MIC is the mean of three biological replicates (black dots) for each strain.

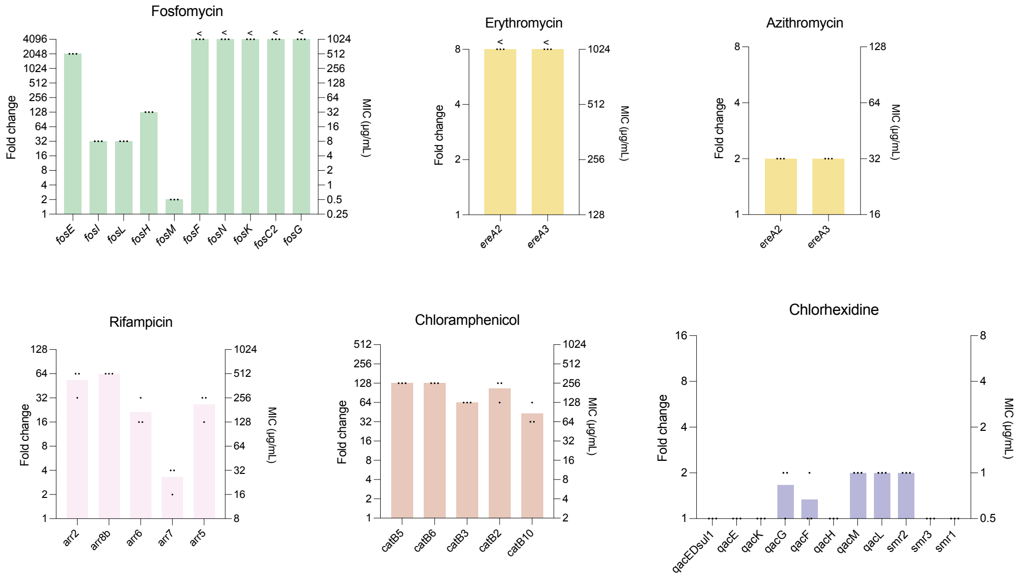


**Supplementary Figure S4. MICs of other ARCs in anaerobiosis.** Antimicrobial resistance to fosfomycin, erythromycin, azithromycin, rifampicin, chloramphenicol, and chlorhexidine is shown as MIC (µg/mL) in the right axis, and resistance fold increase compared to pMBA in the left axis. The MIC is the mean of three biological replicates (black dots) for each strain.

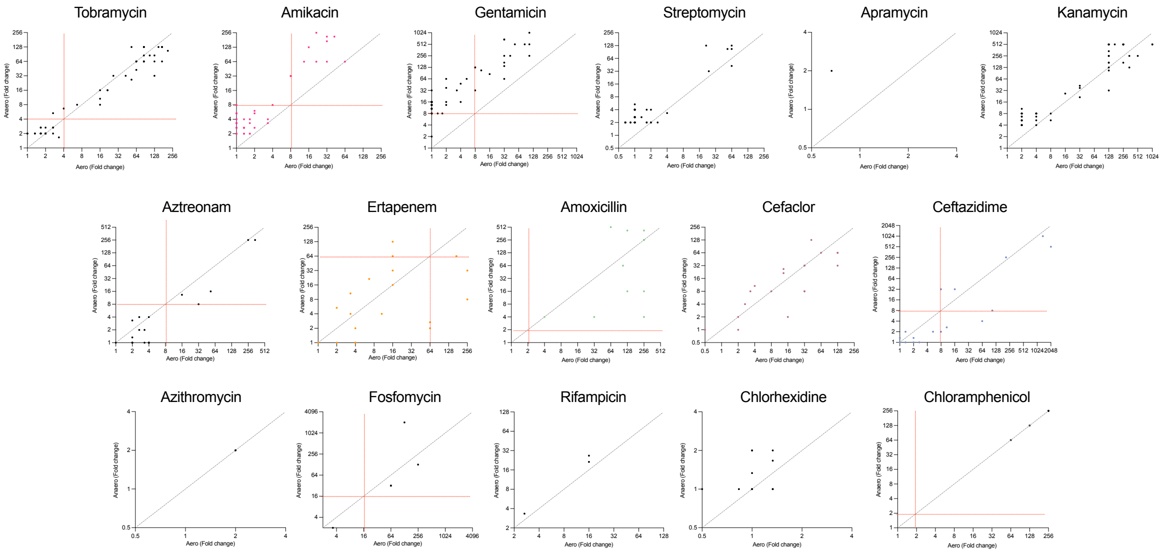


**Supplementary Figure S5. Correlations between antimicrobial effects of each ARC in aerobiosis and anaerobiosis.** Resistance fold change (in comparison with the parental strain pMBA) of each ARC in aerobic and anaerobic conditions is shown in the X and Y axes respectively. Red dotted lines represent the clinical breakpoint (EUCAST) for *E. coli* against each antimicrobial.

A)

B)

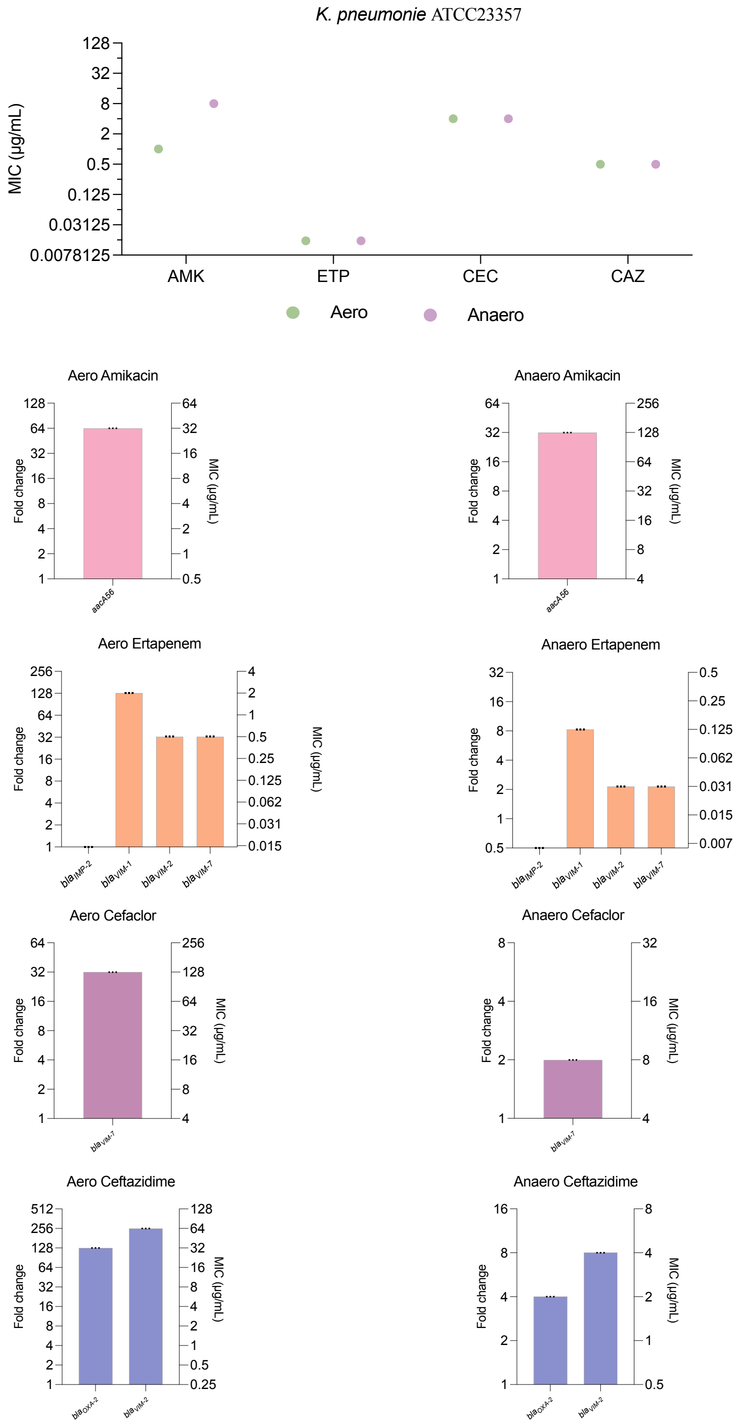


**Supplementary Figure S6. MICs for *K. pneumoniae* ATCC23357. A)** MIC values are shown as the mode of at least three biological replicates. Green dots represent aerobic measures while purple dots represent anaerobic MICs. AMK: amikacin, CEC: cefaclor, CAZ: ceftazidime, ETP: ertapenem. **B)** MIC quantitation of ARCs cloned in *K. pneumoniae* ATCC23357 in aerobic and anaerobic conditions. Antimicrobial resistance to amikacin, ertapenem, cefaclor, and ceftazidime is shown as MIC (µg/mL) in the right axis, and resistance fold increase compared to pMBA in the left axis. The MIC is the mean of three biological replicates (black dots) for each strain
